# A comparative study of structural variant calling strategies using the Alzheimer’s Disease Sequencing Project’s whole genome family data

**DOI:** 10.1101/2022.05.19.492472

**Authors:** John S. Malamon, John J. Farrell, Li Charlie Xia, Beth A. Dombroski, Wan-Ping Lee, Rueben G. Das, Badri N. Vardarajan, Jessica Way, Amanda B. Kuzma, Otto Valladares, Yuk Yee Leung, Allison J. Scanlon, Irving Antonio Barrera Lopez, Jack Brehony, Kim C. Worley, Nancy R. Zhang, Li-San Wang, Lindsay A. Farrer, Gerard D. Schellenberg

## Abstract

**Background:** Reliable detection and accurate genotyping of structural variants (SVs) and insertion/deletions (indels) from whole-genome sequence (WGS) data is a significant challenge. We present a protocol for variant calling, quality control, call merging, sensitivity analysis, *in silico* genotyping, and laboratory validation protocols for generating a high-quality deletion call set from whole genome sequences as part of the Alzheimer’s Disease Sequencing Project (ADSP). This dataset contains 578 individuals from 111 families.

**Methods:** We applied two complementary pipelines (Scalpel and Parliament) for SV/indel calling, break-point refinement, genotyping, and local reassembly to produce a high-quality annotated call set. Sensitivity was measured in sample replicates (N=9) for all callers using *in silico* variant spike-in for a wide range of event sizes. We focused on deletions because these events were more reliably called. To evaluate caller specificity, we developed a novel metric called the D-score that leverages deletion sharing frequencies within and outside of families to rank recurring deletions. Assessment of overall quality across size bins was measured with the kinship coefficient. Individual callers were evaluated for computational cost, performance, sensitivity, and specificity. Quality of calls were evaluated by Sanger sequencing of predicted loss-of-function (LOF) variants, variants near AD candidate genes, and randomly selected genome-wide deletions ranging from 2 to 17,000 bp.

**Results:** We generated a high-quality deletion call set across a wide range of event sizes consisting of 152,301 deletions with an average of 263 per genome. A total of 114 of 146 predicted deletions (78.1%) were validated by Sanger sequencing. Scalpel was more accurate in calling deletions ≤100 bp, whereas for Parliament, sensitivity was improved for deletions > 900 bp. We validated 83.0% (88/106) and 72.5% (37/51) of calls made by Scalpel and Parliament, respectively. Eleven deletions called by both Parliament and Scalpel in the 101-900 bin were tested and all were confirmed by Sanger sequencing.

**Conclusions:** We developed a flexible protocol to assess the quality of deletion detection across a wide range of sizes. We also generated a truth set of Sanger sequencing validated deletions with precise breakpoints covering a wide spectrum of sizes between 1 and 17,000 bp.

## INTRODUCTION

Human genetic variation includes single nucleotide variants (SNVs), small insertions and deletions (indels) less than 150 bp, and structural variants (SVs) greater than 150 bp. SVs can result from deletions, insertions, and rearrangements that include balanced inversions and translocations or unbalanced repeats, insertions and deletions resulting in copy number variation (CNV) (1–3). SV/indels arise as both single and complex events *via* germline and somatic mutations (4) and contribute significantly to genetic diversity and to disease susceptibility (5–11).

A variety of SV/indel types and sizes can be detected using high-throughput short-read whole-genome sequencing (WGS). Multiple large-scale SV detection studies have been performed such as the 1000 Genomes Project (12), the Cancer Genome Atlas project (13,14), Genome of the Netherlands (15), the UK 10K project (16), gnomAD (17) and CCDG(18). However, SV/indel calling using short-read sequence data continues to be challenging. Multiple algorithms and programs (e.g., Breakdancer, CNVnator, DELLY, Genome Analysis Toolkit (GATK: 3.2) Haplotype Caller, Lumpy, Pindel, Scalpel, and SWAN) (19–26) are available, but many factors continue to hinder accurate and comprehensive identification of SV/indels in sequence data. These confounding factors include complex sequence structure, variability in read depth and coverage across the genome, sequencing bias and artifacts, biological contamination, and mapping and alignment errors or artifacts. Also, computational demands can limit the use of some SV/indel calling programs. Furthermore, SV/indel calling in large samples lacks standards for calling procedures, call-set merging, and quality control (QC). These challenges become even more daunting when merging SV/indel calls from samples sequenced at multiple centers that use different sequencing library designs and protocols. The quality and characteristics of sequence data may vary considerably among samples within and across centers and can affect SV/indel calling sensitivity and specificity (27). We present results from analyses of WGS data generated by the Alzheimer’s Disease Sequencing Project (ADSP) for 578 members of 111 families. We focused on deletions because for the programs we examined, deletions were more reliably called than other SV types. Our work showed that no single caller can accurately detect a broad range of deletion sizes. We developed two systematic approaches for evaluating the sensitivity and specificity of different callers and deletions identified from data generated by different platforms and sequencing centers. Finally, we validated a comprehensive strategy for calling, merging, QC, and genotyping deletions that has high sensitivity and minimizes false positive calls.

## METHODS

### Subjects and generation of WGS data

WGS data were obtained from the ADSP, a collaboration between the National Institute on Aging (NIA), National Human Genome Research Institute (NHGRI), and the Alzheimer’s disease research community (**28**). Details of subject selection and WGS data generation and processing are described elsewhere (**28,29**). In brief, the sample included 498 AD cases and 84 cognitively normal elderly controls from 44 non-Hispanic Caucasian and 67 Caribbean Hispanic families. All studies involved were approved by their respective University Institutional Review Boards (IRBs) and the overall study was approved by the University of Pennsylvania IRB. WGS data were generated using Illumina’s 2500 HiSeq platform by the NHGRI’s large-scale sequence and analysis centers (LSACs) at the Baylor College of Medicine (BCM), the Broad Institute (BI), and the McDonnell Genome Institute at Washington University (WashU). BCM provided 166 samples with a mean template size of 370 bp (SD=12.4 bp). For the BI, 232 samples were sequenced with a mean template size of 335 bp (SD=1.4 bp). WU provided 186 samples with three library preparations targeted at insert sizes of 200, 400, and 550 bp. These three library sizes were chosen to increase SV calling accuracy by incorporating longer reads; however, there was considerable size heterogeneity in the 550 bp read group. Three samples from one family were sequenced at all three LSACs as triplicates for evaluating and adjusting for center-specific sequencing effects.

### Deletion Variant Calling Protocol

Two complementary pipelines for deletion calling, merging, genotyping, and reassembly were implemented (Figure 1). In one approach, each genome was divided into 75 regions excluding telomeres and centromeres and called in parallel using Scalpel (**26**) to reduce processing time across the entire genome. Scalpel reassembles gapped alignments using the de Bruijn graph method to increase calling specificity in regions characterized by complex repeat structures. Scalpel was also used to generate precise breakpoints *via* local assembly within a 1,000 bp capture window for the whole genome. GenomeSTRiP (**30**) was used to perform joint-genotyping to provide missing genotype information and further refine calls. The second deletion calling pipeline was based on Parliament (**31**), which creates a unified project-level variant call file (pVCF) by combining and filtering calls based on consensus and quality metrics from eight indel/SV callers including Scalpel (Table 1). Parliament also provided gene annotation, genotyping, and local hybrid assembly. Because Parliament is computationally intensive, we limited the analysis to deletions > 200 bp. The functional annotation of each variant was determined using SNPeff (**32**).

**Figure 1.**
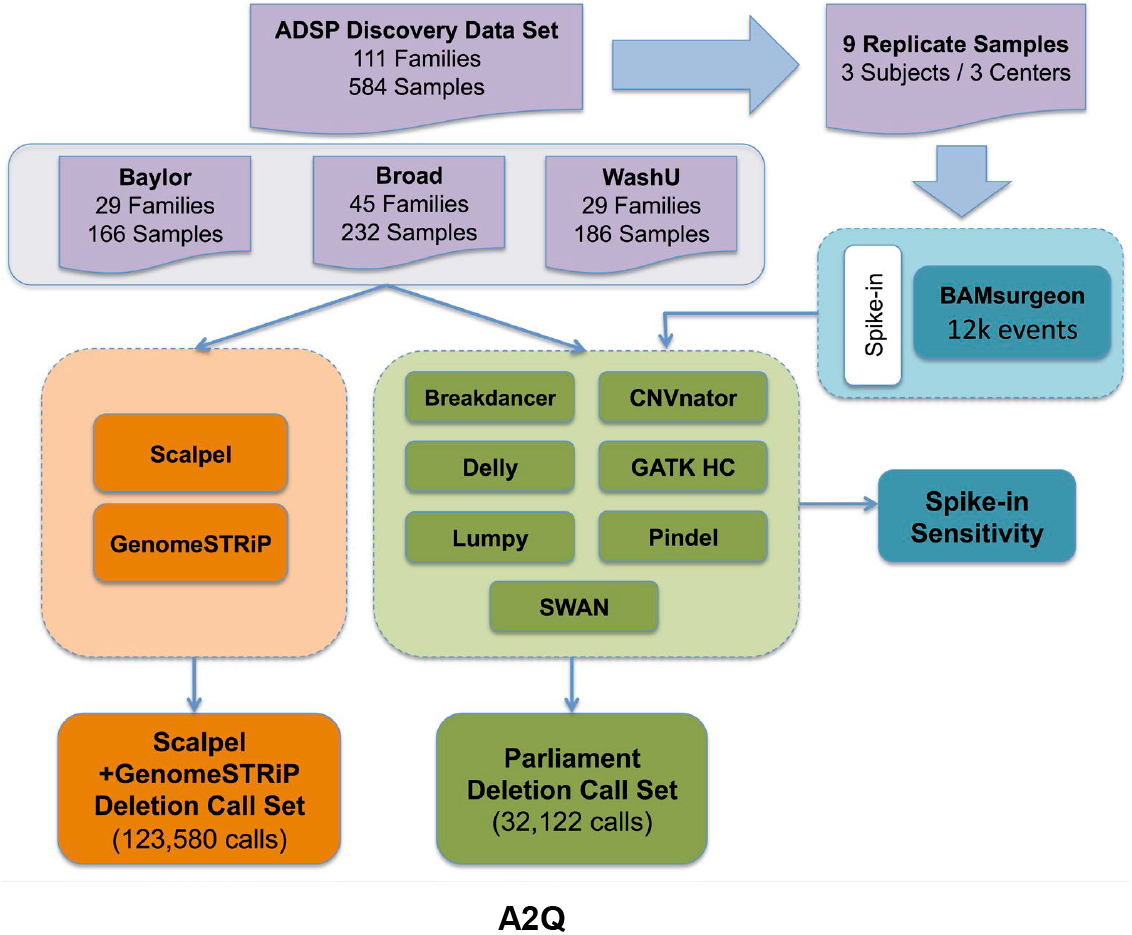
Overview of ADSP’s SV/indel calling and analysis pipeline. Two parallel pipelines, Scalpel+GenomeSTRiP (orange) and Parliament (green), were combined to perform SV/indel call merging, QC, genotyping, and re-assembly for 584 samples from three sequencing centers. Nine replicated samples were used to measure individual SV/indel caller sensitivity *via* variant spike-in studies.

**Table 1.** Overview of SV/indel callers evaluated. Column 1 provides the caller evaluated. The second column provides the software version used for each caller. ‘Sequence Feature’ provides the method used to determine events such as read-pair (RP), split-read (SR), read-depth (RD), soft-clip (SC), and local assembly (AS). Columns 5 and 6 provide whether each caller supplies precise genotypes and breakpoints.

| Caller | Software Version | Sequence Feature | Calling Range (bp) | Exact Genotype | Precise Breakpoint |
| --- | --- | --- | --- | --- | --- |
| Breakdancer | 1.1 | RP | 1 – 10,000 | Yes | No |
| CNVnator | 0.3 | RD | 200 – 10,000 | No | No |
| DELLY | 0.5.6 | RP and SR | 15 – 10,000 | Yes | Partial |
| GATK HC | 3.2 | RP | 2 – 300 | Yes | Yes |
| LUMPY | 0.2.10 | RP, SR and RD | 1 – 10,000 | No | No |
| PINDEL | 0.2.5a3 | RP and SR | 1 – 10,000 | Yes | No |
| Scalpel | 0.5.3 | AS | 1 – 1,000 | Yes | Yes |
| SWAN | 0.3.0 | RP, SR, and SC | 1 – 10,000 | No | Partial |

### Sensitivity Analysis Using Simulated Spike-in Data

We estimated sensitivity by “spiking-in” SV/indels using BAMSurgeon (**33**) into triplicated samples (three samples sequenced at all three LSACs). First, we generated a list of predefined SV/indels, including 4,040 deletions and insertions and 1,560 inversions and tandem duplications, totaling 11,200 events. SV/indels ranged in size from 2 to 5,000 bp and were spiked into all autosomes for the three sample replicates (nine files total). Half of the events were inserted as heterozygotes and half as homozygotes. BAMSurgeon failed to add in a small fraction (2.92%) of the attempted events, and those sites were excluded from sensitivity analysis. For sites where BAMSurgeon succeeded, there were minor discrepancies in the exact breakpoints of the actual spike-in as compared to its targeted location. These minor breakpoint discrepancies did not affect the results because we applied a 50% reciprocal overlap for detecting spiked-in events. Finally, SV/indels were called for the nine spiked-in samples to measure the sensitivity of each caller across the full range of sizes. Since the true events were known or spiked-in, the sensitivity (*Eq. 1*) of each SV/indel caller was estimated as:

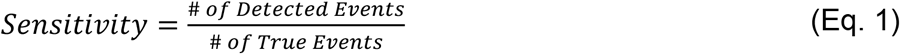

### D-score: A metric for evaluating SV/indel caller specificity in family studies

To ascertain the specificity of deletion calls, we developed the following family-based metric called the deletion or D-score:

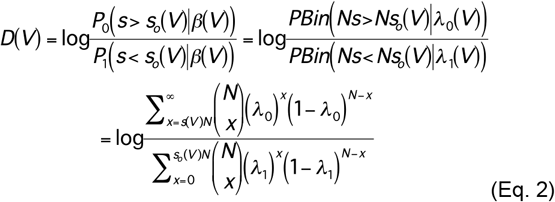

*s_o_*(*V*) is the observed sibling sharing frequency for a variant (*V*) and 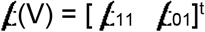 is the caller sensitivity vector of detecting homozygous and heterozygous variants of the same type and size as *V*. Caller sensitivity vectors, 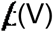, were calculated from the spike-in study results for each caller. *s* represents the sharing frequency across all unrelated subjects. *P_0_* and *P_1_* represent the probability of observing the null and alternative hypothesis, respectively. *D(V*) represents the log likelihood ratio that compares the probability of observing *f_0_*(*V*) under the assumptions that H_0_: *V* is false or H_1_: *V* is true. For example, when *s* (unrelated sharing frequency) is greater than *s_o_*(*V*) (sharing frequency among siblings) the null hypothesis is more likely to be supported.

The allele frequency was computed for *V* in siblings (*f*_1_) and in all samples (*f*_0_) based on the estimated caller sensitivity for each variant by the following

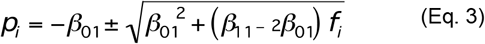

Only one of the roots of *p_i_* will satisfy the requirement 0<*p_i_*<1 and was used to calculate the mean expected sibling sharing for the two hypotheses using the following equation:

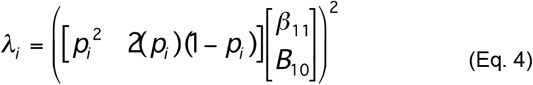

The theoretical range of the D-score in this dataset was between −40 and 40. The D-score metric does not require deletion genotype information and therefore can be used to evaluate caller specificity in the absence of genotyped calls, as is the case with many SV/indel callers.

### Kinship coefficient

To assess overall call set quality, a kinship coefficient was calculated using KING (34) for all sibling pairs with genotype information. Because a kinship coefficient of 0.25 is expected for the pooled set of heterozygous joint-genotyped calls, departure from this value indicates systematic errors in SV/indel calling. Because multigenerational data are usually not available in family studies of AD, the kinship coefficient has greater utility than a check for mendelian inconsistencies and is useful for measuring the overall quality of the genotypes. This metric is analogous to the Ts/Tv ratio for SNVs which has an expected value of 2.15 for high quality SNV data sets.

### Quality control

Many false positives are the result of poor mapping quality between two or more sites and are characterized by excess heterozygosity. Therefore, a Hardy-Weinberg equilibrium p-value threshold of 5×10^−8^ was applied to filter calls with excess heterozygosity. The BLAST-like Alignment Tool (BLAT) (**35**) was used to filter deletions with a low predicted mapping quality or that map to many sites (N>100) in the genome. Finally, deletions with an alternate allele count of less than five were removed from the final call set. Parliament’s consensus and QC strategy proved to be useful in improving call quality by combining call set metrics and applying heuristics to reduce false positives.

### Computational performance of SV/indel callers

Computational performance benchmarks were obtained for the eight SV/indel programs based on analysis of 20 randomly selected subjects. Performance benchmarks were derived using automated scripts and include total run time, peak central processing unit (CPU) usage, peak memory usage, and processing core hours. All data were processed using an © Amazon’s Elastic Cloud 2 (EC2) extra-large instance with © Intel © Xeon 2.4 GHz CPUs. Scalpel benchmarking results were excluded from this analysis due to its extreme computational demands for processing WGS data.

### Laboratory validation of deletion calls

Subsets of Scalpel and Parliament-derived deletions of different sizes were selected for validation based on three methods: randomly selected events within specified size bins, predicted LOF, and proximity to 74 candidate AD loci with strong genome-wide association signals. These candidate AD loci were curated from GWAS, candidate gene studies, and multiple family-based studies (36–66). Validation was performed by polymerase chain reaction (PCR) across the deletion with custom designed primers followed by Sanger sequencing. For the Scalpel-derived deletions, the variants were binned by base pair length (2-19 bp, 20-40 bp, 41-60 bp, 61-80 bp, 81-100 bp, and 101-900 bp). The size ranges examined for the Parliament-derived deletions were 101-900 bp, 901-1,000 bp, and 1,001-17,000 bp. The BLAST-like alignment tool (BLAT) from University of California, Santa-Cruz (UCSC) Genome Browser (35) was used to search and align variant sequences and surrounding sequences to the human genome. Because BLAT has a minimum requirement of 20 bp, sequences smaller than 20 bp were queried by adding flanking sequences upstream and downstream of the test sequence to bring the length up to 20 bp. Both UCSC HG19 and HG38 reference genomes were queried using BLAT. Additionally, for each deletion, 100 bp sequences flanking either side of the event were also queried against BLAT as a contiguous 200 bp sequence (*i.e*., variant deletion sequence removed). BLAT alignment allowed visualization of the deletion and surrounding sequence in terms of proximity to genes and repeat sequence and facilitated the identification of instances of clear mis-mapping. Sequence surrounding the variants was extracted from HG38 and used for primer design. For variants where a PCR product of ≤1,200 bp was expected (including the variant sequence), primers were designed outside of the breakpoints to amplify across the deletion sequence. For deletions where the reference allele was too large to be amplified by a 1,200 bp PCR product, a double PCR approach was used. For the first PCR, one primer was designed within the putative deletion sequence while the other primer was placed external to the deletion breakpoint. Samples containing the reference allele and not containing a deletion would give a product with this PCR. For the second PCR, both primers flanked the putative deletion. Only samples, which contained the deletion, would yield a product for this PCR. Samples from three individuals reported as heterozygous or homozygous deletions were used for sequence validation as well as one control (or reference) sample. When possible, samples from multiple families were used for validation.

## RESULTS

We generated deletion calls for the ADSP Discovery Phase WGS using 8 different programs (GATK haplotype caller, Scalpel, Breakdancer, CNVnator, Lumpy, Pindel, Swan, and DELLY)(Figure. 1). These programs use different sequence features and analyze different event sizes (Tables 1 and 2). To determine the properties of the data generated by each program, we systematically evaluated sensitivity and specificity. Because the sequence data was generated at three different LSAC sites using libraries with different characteristics, we evaluated data from each site. We also benchmarked the computational resources needed for each program.

**Table 2.** Total calls by eight SV/indel callers. Number of pre-QC calls for all eight callers.

| Caller | Number of Calls |
| --- | --- |
| Breakdancer | 3,484,082 |
| CNVnator | 2,572,070 |
| DELLY | 1,685,852 |
| GATK Haplotype Caller | 232,366 |
| LUMPY | 1,003,953 |
| PINDEL | 2,613,604 |
| SWAN | 1,945,009 |
| Scalpel | 1,441,659 |
| Total | 14,709,212 |

### Sensitivity Analysis

Sensitivity was evaluated by inserting deletions and insertions into WGS data generated at each LSAC. Sensitivity for detecting the inserted deletions varied among callers and, to a lesser extent, the source of the sequence data, and was dependent on the size of the deletion (Figure 2). For short deletions (30-500 bp), Scalpel showed the best sensitivity (~85%) and was closely followed by Pindel. Pindel showed good sensitivity up to 1000 bp. GATK-haplotype caller showed a sensitivity of ~75% for events up to 100 bp but fell off rapidly above this size range. For larger events, Lumpy and SWAN both showed good performance up to 5,000 bp with SWAN able to detect even larger events. DELLY showed reasonable sensitivity in the 500 to 5,000 bp range but when compared to other programs, results were more influenced by the source of the data. For example, DELLY had lower sensitivity when calling genomes sequenced by BCM in the 200-500 bp bin as compared to those from WashU and BI. SWAN was the most sensitive caller across all sizes and sequencing centers, perhaps because it accounts for various sequencing characteristics such as multiple insert-size libraries and soft-clipped reads (**24,67**). CNVnator and Breakdancer showed poor sensitivity for all size ranges. Our results show that sensitivity varies among callers and for different size ranges but is relatively insensitive to the sequencing site.

**Figure 2.**
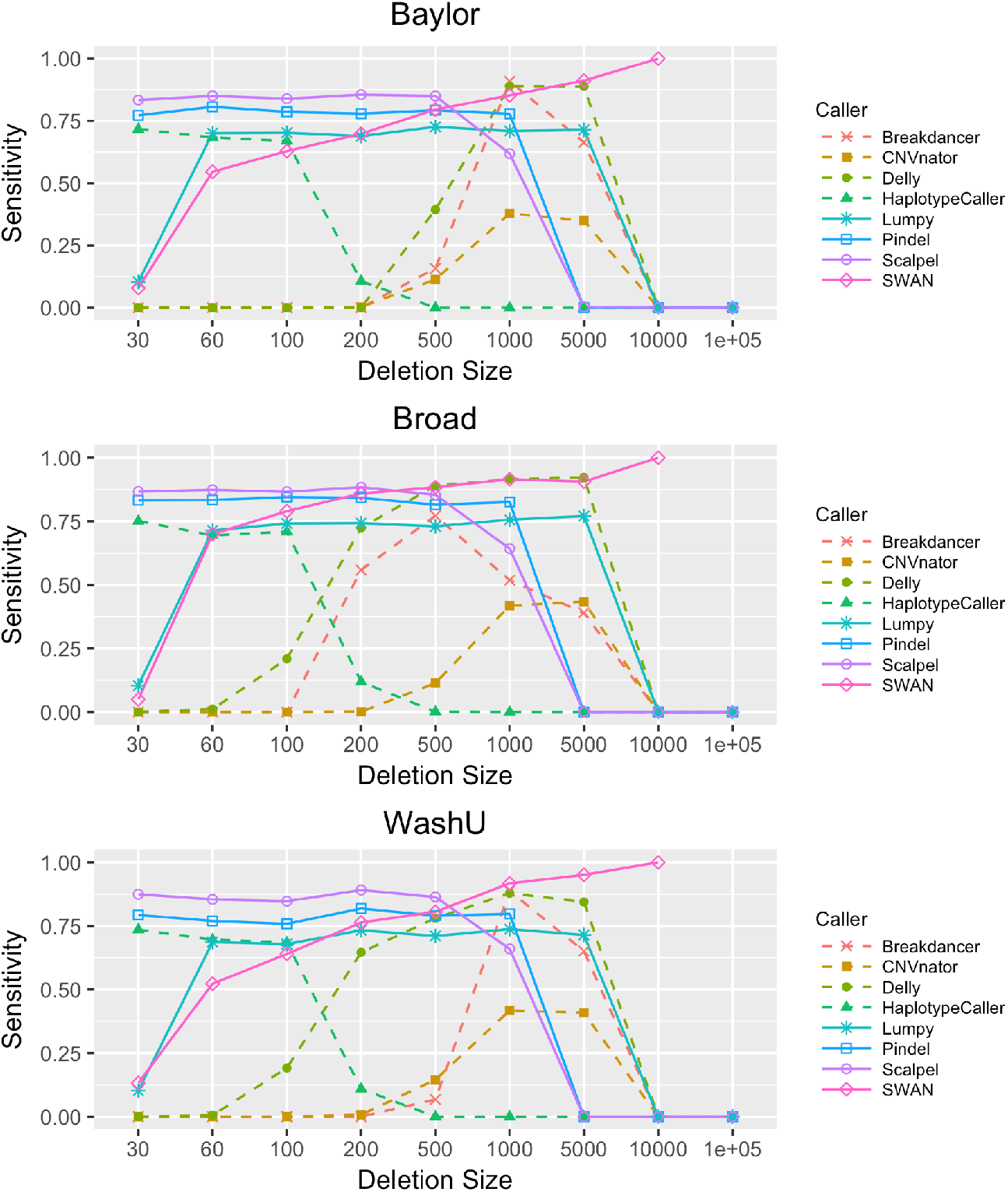
SV/indel caller sensitivity stratified by sequencing center and caller. Sensitivity rates were derived for all eight callers using *in silico* variant spike-in on nine sample replicates. Sensitivity is provided for all three centers (Baylor, Broad, and WashU). Biological replicates are three individuals in one family that were sequenced at the three centers. Sensitivity rates are provided across a large range of event sizes [30 bp-10 kb]. Sensitivity rates are largely consistent across centers.

### Specificity analysis

We assessed caller specificity using the D-score method. LUMPY was the best performing program with D-scores between 5 and 10 for deletions from 30 to 10,000 bp (Figure 3). The results were independent of the sequencing center. Scalpel also yielded highly specific calls, particularly in the 200 to 1,000 bp range with D-scores ranging from 5-8. Median D-scores for deletion calls from SWAN, Pindel and Breakdancer were between 3 and 5, but the results were dependent on the sequencing center. Other programs yielded calls with lower specificity that were greatly influenced by sequence source. We also applied the kinship coefficient to evaluate and calibrate the quality of deletion calls and measure the impact of QC steps on call specificity (Figure 4). Prior to data cleaning, the kinship coefficient was much greater than the expected value of 0.25 for siblings for events ranging from 21-350 bp. After removing deletions showing excess heterozygosity, the kinship coefficient of the Scalpel genotypes approached 0.25 for all deletion sizes (Figure 4a). Comparison of kinship coefficient metrics also showed that the quality of GATK Haplotype calls decreased as the deletion size increased and the coefficient was 0 at >50 bp. In contrast, a kinship coefficient of 0.25 was maintained for Scalpel calls for deletion sizes between 20 bp and 400 bp, showing that the Scalpel calls are more reliable in this size range. This work shows that the specificity of calls from different programs varies depending on the size of the event detected and can be influenced by the source of the sequence data.

**Figure 3.**
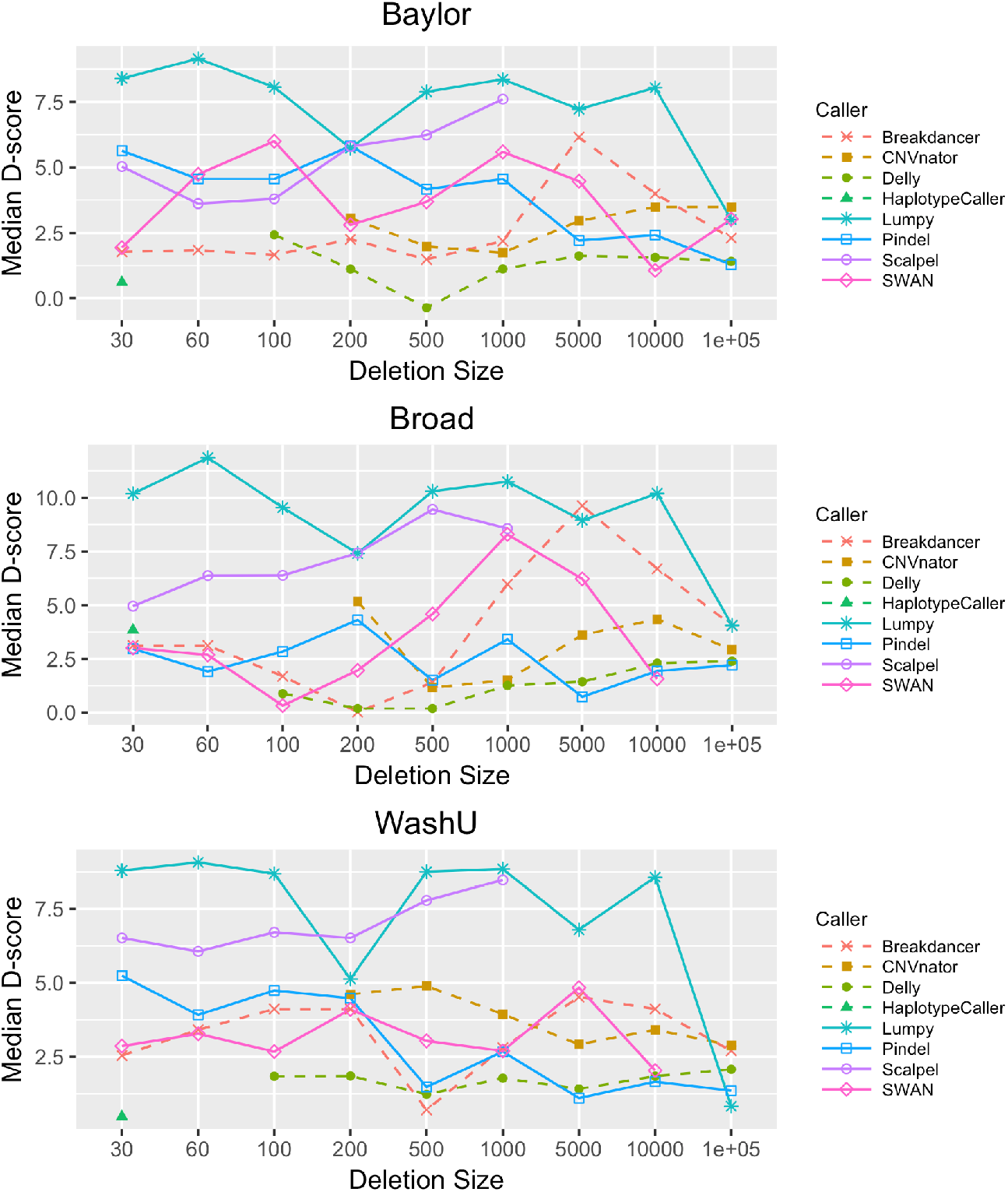
SV/indel caller specificity using the D-score stratified by sequencing center and caller. Specificity rates are provided for all eight callers from 30 bp to 10 kbp using our D-score method. D-scores were calculated for each of three sequencing centers (Baylor, Broad, and WashU). D-scores are quite consistent across centers.

**Figure 4.**
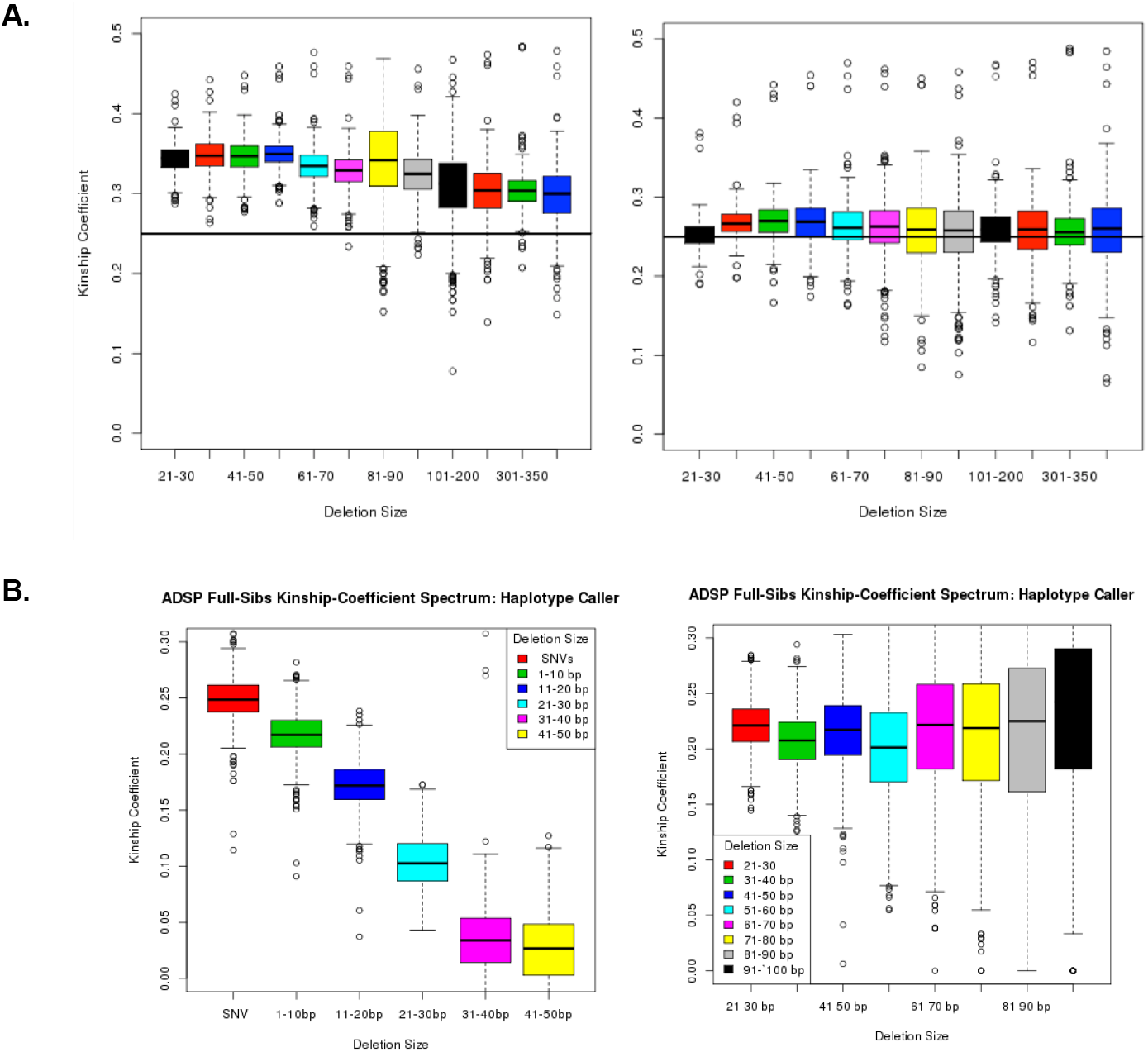
Kinship coefficient by deletion size. **A. pre- and post-QC.** Kinship coefficients for pre- (left) and post-QC (right) calls ranging from 20-400 bp were calculated for all sibling pairs. QC filtering to reduce excess heterozygosity resulted in coefficients that approximated the expected value of 0.25. **B.** Kinship coefficients for GATK Haplotype Caller (left) and Scalpel (right) were calculated using all sib pairs. The coefficient is 0.25 for SNVs called by the GATK Haplotype Caller but declines progressively to 0 with increasing deletion sizes. In contrast, the coefficient approximated 0.25 for Scalpel calls across all SV/indel bin sizes.

### Assessment of SV/indel caller computational requirements

We measured computational performance metrics for seven of the eight callers used in this study (Figure 5, Supplementary. Table 1). Scalpel was excluded from performance benchmarking due to its extreme CPU demands and total runtime. To generate these benchmark metrics, we processed 10 BAM files (mean size of 209.05 MB) from the ADSP’s discovery (disc) phase and 10 BAM files (mean size of 54.58 MB) from the discovery extension (disc+ext) phase. Among the tested callers, SWAN had the highest memory demands and required more than 10-times greater run time compared to other programs. Breakdancer was the second longest running SV caller evaluated. DELLY, Lumpy, GATK, and SWAN all had similar CPU demands. While Scalpel and SWAN ranked high in terms of sensitivity and specific, the run time computational requirements preclude the use of these programs on large datasets.

**Figure 5.**
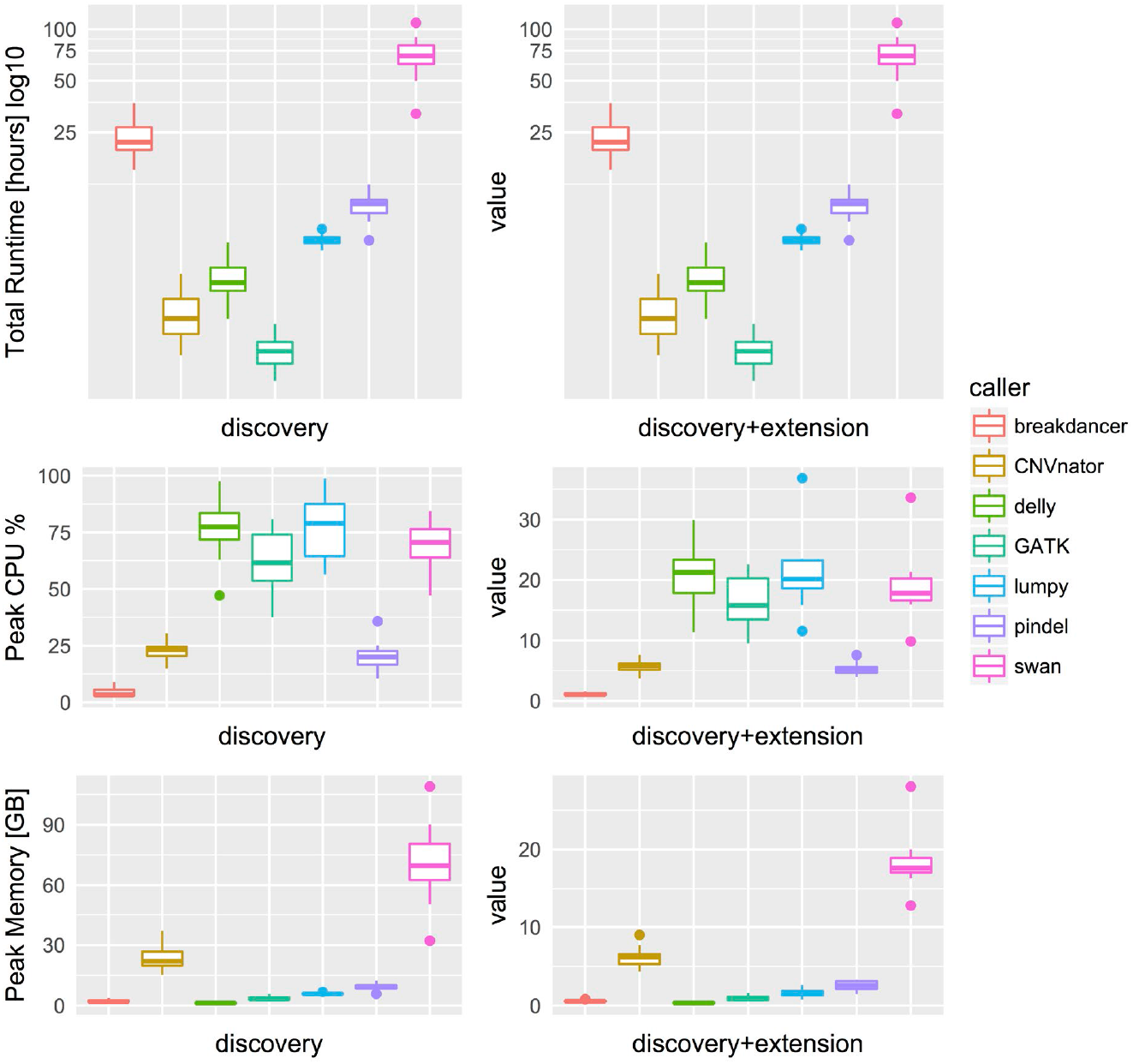
Three performance metrics for seven SV/indel callers. Top row provides total runtime in hours, middle row provides peak CPU percentage, and bottom row provides peak memory in gigabytes for 10 discovery phase (left column) and 10 discovery-extension phase (right column) samples.

### Generating an ADSP deletion call set

All 584 samples were called in parallel *via* two independent production pipelines, Scalpel+GenomeSTRiP and the Parliament toolkit (Figure 1). Given that the sensitivity and quality for the GATK haplotype caller dropped off significantly with deletions of size greater than 20 bp, the pipelines focused on deletions greater than that size range. Localized assembly and break-point refinement on gapped alignments was performed with Scalpel to increase calling accuracy of deletions as large as approximately 900 bp. Of the 123,581 deletions detected by Scalpel and genotyped with GenomeSTRiP, 100,678 sites remained after removal of excess heterozygotes (N=17,286), homozygous reference (N=5,014), and call rate <less than 90% (N=603). The number of deletions called dropped off exponentially as the deletion size increased except for a spike in the number of deletions related to Alu retrotransposons (Figure 6). The frequency of these events peaked around 350 bp which corresponds to lengths of Alu transposons, and this size distribution is expected and consistent with that observed in other studies(17). The Parliament pipeline genotyped more than 14 million SVs from the eight callers listed in Table 2. The mean number of calls per program was slightly greater than 1.8 million. Due to computational requirements, the sites genotyped were limited to those greater than or equal to 100bp. A total of 32,122 remained post-QC and the size distribution of these calls shows the Alu peak at ~350 bp (Figure 7). The distribution of functional annotations of these variants is shown in Table 3. Comparison of the deletion calls generated by the two pipelines in the size ranges that overlapped (100-900 bp) identified 3,401 deletions (mean size = 330 bp, range 207 - 620 bp) that shared a base location for at least one breakpoint (Supplementary Figure 1) in the size bin with deletions common to both callers.

**Figure 6.**
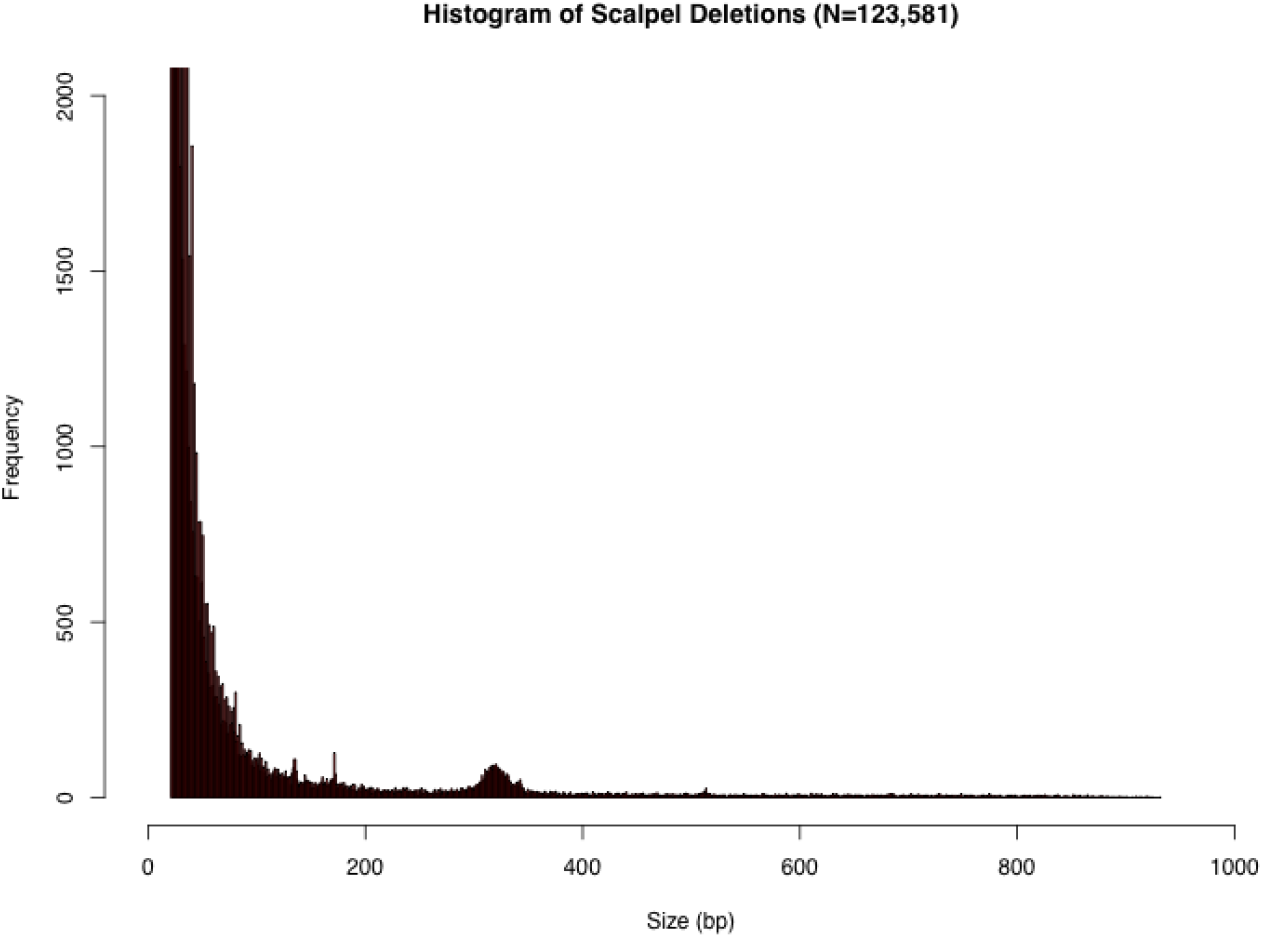
Histogram of Scalpel deletions by size. All Scalpel deletions (N=123,581), ranging from 20 to 900 bp. Y-axis is truncated at 2,000 calls. Alu peak is seen near 350 bp.

**Figure 7.**
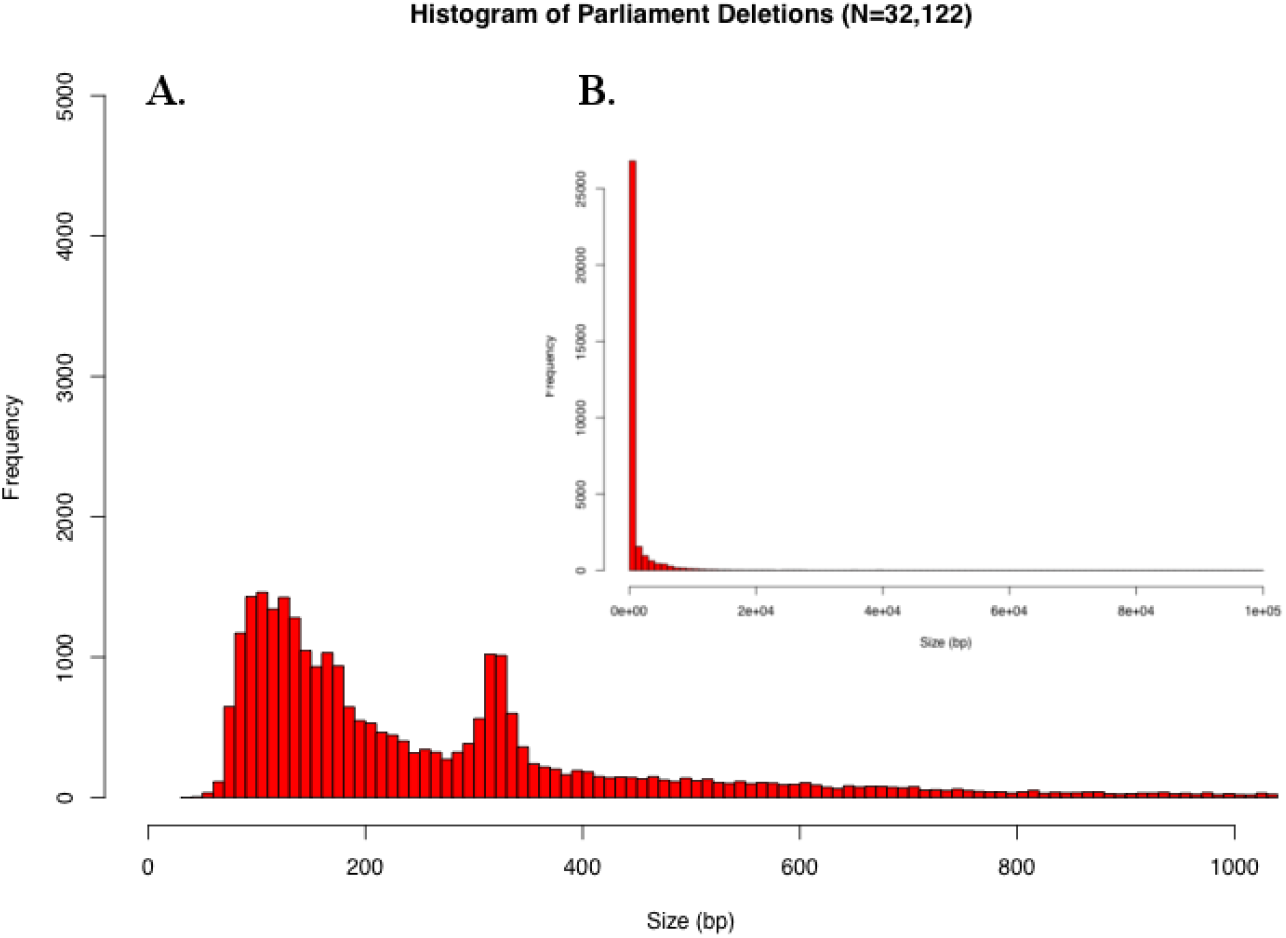
Histograms of Parliament deletion frequencies by size. A) Bottom-left: Histogram of Parliament deletions (N=32,122) ranging from 20 to 1,000 bp. Alu peak is seen at ~350 bp; and B) Upper-right: Full histogram of all Parliament calls (N=32,122) ranging from 1-10,000 bp.

**Table 3.** SNPEff functional annotation categories for Scalpel and Parliament calls. Breakdown of genomic functional annotation terms provided by SNPEff. There are slightly more annotation terms than loci as some loci overlap more than one region.

| Functional Annotation Term | Scalpel | Parliament | Parliament + Scalpel |
| --- | --- | --- | --- |
| Intergenic | 59,595 | 15,238 | 1,900 |
| Coding | 51,849 | 14,724 | 28 |
| Splice site | 339 | 959 | 0 |
| Intronic | 492 | 151 | 1,475 |
| 5 prime UTR | 201 | 113 | 0 |
| 3 prime UTR | 739 | 201 | 0 |
| Other | 10,857 | 887 | 0 |
| Totals | 124,072 | 32,273 | 3,403 |

### Laboratory Validation of Deletion Calls

To validate deletion calls, we performed Sanger sequencing on putative deletions. We sequenced 106 deletions called by Scalpel ranging in size from 2 – 900 bp (Table 4 and Supplementary Table 3). When smaller deletions were randomly selected, 87.5% of events between 2 and 100 bp were validated by Sanger sequencing (100% of the events under 20 bp and 80% of events between 80-100 bp were confirmed by Sanger sequencing). For loss of function deletions and those near AD genes (+/- 500 kb, Supplementary Table 2) in this size range, slightly higher validation rates were observed (average 93% and 95%, respectively). For randomly selected large events (between 101-900 bp), the validation rate fell to 17%. However, when large SV/indel calls were pre-screened to remove deletion sequences found at multiple regions of the genome, the validation rate increased to 50%. Deletions near AD genes and LOF variants had a higher validation rate (83% and 75%, respectively).

**Table 4.**
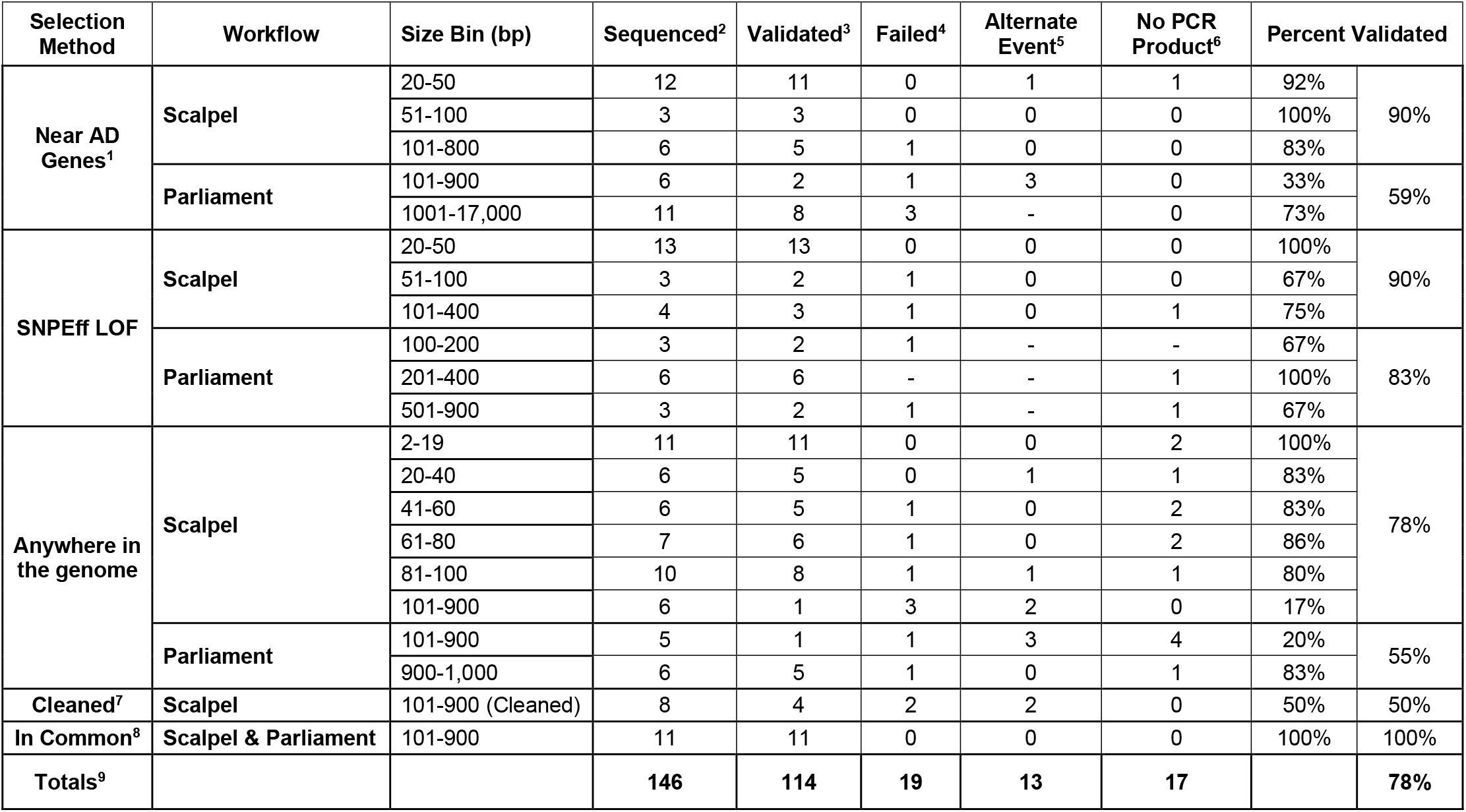
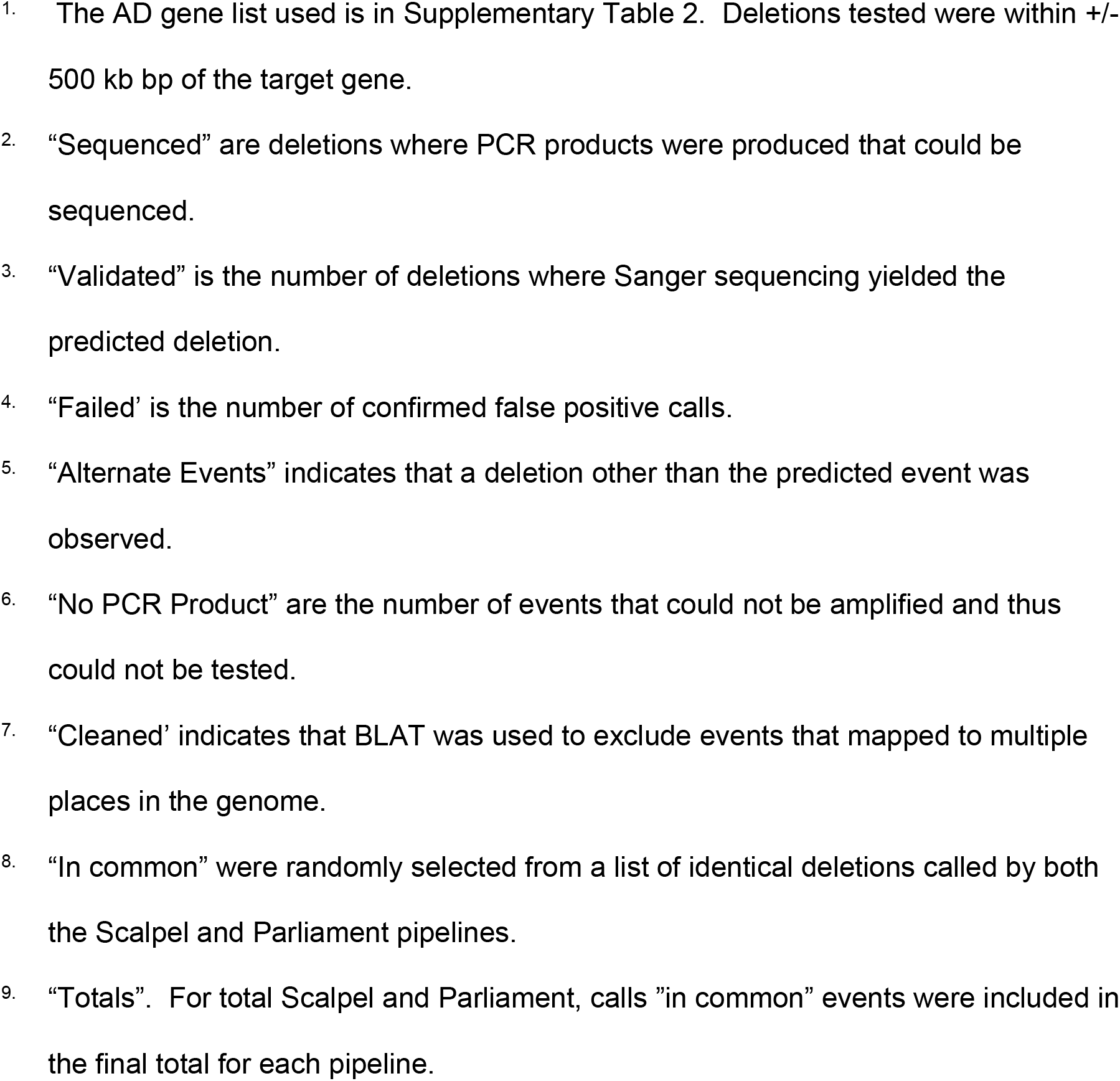
Deletion validation results.

For Parliament pipeline calls, 20% of randomly selected deletions in the 101-900 size range could be validated. For Parliament calls near AD genes and LOF variants, calls were validated at a higher rate (33% and 83%, respectively). For larger SVs calls, the validation rate ranged from 73% - 83%. When we examined calls made by both Parliament and Scalpel, all deletions tested (n = 11) could be validated. The mean D-score for validated deletions (8.12, sd = 10.98, n = 114) was significantly greater than the mean for deletions that were not validated (2.52, sd = 4.98, n = 19, p = 0.0075). This Sanger sequencing validation of deletions demonstrates that the variants called by Scalpel, particularly within the 2-100 bp size range, are highly reliable and are suitable for genetic association studies.

### Deletions near AD genes

To detect possible AD pathogenic variants, we looked for deletions in a +/- 500 kb window bracketing candidate AD genes, focusing on deletions in gene functional units (coding regions, 5’ and 3’UTRs, promoters, and splice junctions). This window was selected to capture genes regulated by cis-acting elements impacted by peak GWAS variants that influence expression of causal AD genes. We identified deletions in the vicinity of 24 AD candidate genes (Supplementary Table 5) that could be validated by Sanger sequencing. One pathogenic deletion identified using Scalpel was a 44 bp deletion in exon 14 of *ABCA7* (rs142076058, p.Arg578 fs). Subsequent work in a larger sample showed that the deletion was associated with AD in African American populations (68). For the remaining confirmed SVs, we tested the segregation of the SVs in the families by requiring that at least 75% of the patients with LOAD and WGS data in the families were carriers. We found segregation of six SVs in at least one family near *IQCK, FBXL7, INPP5D, SPDYE3*, and *SERPINB1* (Supplementary Table 6). A 21 bp coding deletion was identified (rs527464858) in *GIGYF2*, a gene that encodes GRB10 interacting GYF protein 2. This protein regulates tyrosine kinase receptor signaling. The *GIGYF2* deletion is in an imperfect CAG repeat sequence and is ~270 kb from rs10933431, the top SNV for *INPP5D* (P = 3.4 × 10^−9^, OR = 0.91 CI 0.88-0.97) (36). This deletion was observed in 46 cases and 3 controls in both NHW and CH populations. Note that in our study, there were more cases (n = 498) than controls (n = 86) and some of these subjects are related (n = 111 families). This deletion was observed in 10 CH and 13 NHW families. Co-segregation showed that the variant segregated with AD status in three NHW families and one Hispanic family. A number of studies found variants in *GIGYF2* potentially associated with an autosomal dominant form of Parkinson’s disease (69–71), particularly in European populations but not in Asian cohorts (69–72). While several SNVs in *GIGYF2* may be associated with PD, most studies do not confirm an association of this gene with PD (71), and a large meta-analysis did not find that PD was associated with the poly Q region deletion described here (72).

## DISCUSSION

We developed novel approaches for detecting deletions and evaluating them for sensitivity, specificity, and validity. These methods were applied to WGS data obtained from 578 participants of the ADSP. We evaluated eight SV/indel callers on data generated at three sequencing centers, each of which generated sequence libraries using different protocols. Although sequencing library heterogeneity did not appreciably influence results obtained with most programs, call validity (deletion detection by an orthogonal method) varied by size and calling program. Our results revealed that no single calling program could reliably and accurately detect deletions in all size rangers. Ultimately, we effectively detected and genotyped deletions in the WGS dataset using a combination of SV/indel callers, applying several QC filters, and validating calls by Sanger sequencing.

We evaluated the sensitivity of multiple SV/indel callers by *in silico* insertion of deletions and insertions into ADSP biologic replicate sequence data. This simulation exercise suggests that Scalpel has the highest sensitivity for deletions in the 30-500 bp range. The sensitivity performance was closely matched by the programs Pindel and GATK Haplotype caller, but the latter only for smaller events. Also, the specificity of calls by Pindel was much less than Scalpel because of the excess number of events called by this program. We measured specificity of the SV/indel programs using the D-score, a measure that compares deletion sharing between related and unrelated individuals, and the kinship coefficient which allows a comparison of the observed number of deletion calls with the number of expected calls among individuals with a defined degree of relationship. Scalpel and Lumpy showed the best specificity across a broad size range from 30 – 1,000 bp and were relatively insensitive to sequence library differences. In contrast, output from other callers was more sensitive to the source of the sequence data.

We developed a comprehensive pipeline for calling, merging, QC, genotyping, and break-point refinement of deletions using Scalpel and GenomeSTRIP (Fig. 1). As expected, the most common deletions were small and we observed an excess of deletions of approximately 350 bp in length, many of which are likely Alu repeat sequences (Figs 5 and 6). For the size bin 20-100 bp, Sanger sequencing validated more than 87.5% of randomly selected deletions and 90.1% of all deletions (random, near AD genes, LOF variants) (Table 4). This size bracket included 82,180 deletions and accounts for 88.7% of all deletions detected by Scalpel (n = 92,659 total deletions, Supplementary Table 4). In addition, the Scalpel dataset had a kinship coefficient near the expected value of 0.25 for siblings after the removal of sites with excess heterozygosity.

Our study has several noteworthy strengths. First, we developed methods for evaluating deletion specificity in family-based studies (D score). This allowed us to directly compare different methods of deletion calling directly using study sequence data. Also, the D-score can be used to prioritize SVs for targeted validation. Second, we used the kinship coefficient metric as a method to measure the overall quality of the call set genotypes and evaluate quality control measures applied to family-based data. Third, we generated spiked-in data sets that allowed for the evaluation of sensitivity in the sequence data used in this study. Fourth, an orthogonal method (Sanger sequencing) was used to validate candidate deletions and to identify characteristics of true calls. Fifth, the high-quality deletion calls from Scalpel, particularly those under 100 bp, can be used as a gold standard for comparison with calls from other programs that are computationally less intensive. Sixth, we cataloged deletion sites with precise breakpoints that can be directly genotyped in WGS CRAMS with other genotyping tools such as Graphtyper and Paragraph. Last, we detected a deletion in *ABCA7* that was subsequently shown to be pathogenic. This illustrates the validity of our approach for identifying AD-related deletions..

Our conclusions and recommendations for deletion calling have some limitations. Although the D-score and kinship coefficient are useful specificity measures, they require family-based data. Also, because the D-score method relies on comparison of the deletion frequency in the general population (i.e., unrelated individuals) *versus* related individuals, it does not perform well for deletions that are very rare (less than 20 instances in a data set) or very common with allele frequencies approaching 50%. A minimum of two SVs are needed to compute a D-score. In both cases, the resulting D-score will be close to zero. Second, computational requirements need to be considered. Scalpel, while yielding high-quality calls, is not practical when applied to WGS data sets containing more than a few thousand subjects because this program is computationally intensive. However, the Scalpel calls generated here can be used as a benchmark for evaluating sensitivity and specificity of other programs such as more recent versions of GATK haplotype caller (unpublished data). The utility of callers with longer runtimes can be improved by splitting larger chromosomes and processing them in parallel. However, the cost of using some programs such as Scalpel and SWAN may be prohibitive when applied to datasets much larger than the one used in this study. Another limitation of our study is that for associations of deletions with AD, our study is underpowered. Thus we can nominate deletions as candidate pathogenic variants (e.g. Supplementary Table 5) but will need larger follow up studies to confirm true associations (*e.g*. the *ABCA7* deletion). Finally, we only evaluated deletions in this study due to the poor performance of the callers used to detecting insertions and other types of events. Future studies will use other programs that better detect insertions, rearrangements, and copy-number changes.

Findings from this study have multiple important implications. Small deletions represent a substantial portion of genetic variation.(17,73) Larger deletions are rarer and account for a small fraction of total genetic variability but are more likely to be deleterious because they may involve large portions of one or more genes. Given the challenges of accurate SV/indel detection and genotyping, SV/indels larger than a few base pairs are typically not included in genetic association studies. Accurately called and genotyped indels/SVs can increase the scope of both hypothesis-driven and genome-wide association studies. Moreover, similar to single nucleotide variants, SV/indels in the context of a large WGS or WES dataset can be imputed reliably into GWAS datasets derived from SNP arrays. Studies of SV/indels in the future will likely increase and improve our understanding of the genetic architecture of many diseases as more reliable and efficient calling algorithms are developed and validated.

## Acknowledgements

The Alzheimer’s Disease Sequencing Project (ADSP) data used is available through the NIA Genetics of Alzheimer’s Disease Data Storage Site (NIAGADS), data set NG00067. The ADSP is comprised of two Alzheimer’s Disease (AD) genetics consortia and three National Human Genome Research Institute (NHGRI) funded Large Scale Sequencing and Analysis Centers (LSAC). The two AD genetics consortia are the Alzheimer’s Disease Genetics Consortium (ADGC) funded by NIA (U01 AG032984), and the Cohorts for Heart and Aging Research in Genomic Epidemiology (CHARGE) funded by NIA (R01 AG033193), the National Heart, Lung, and Blood Institute (NHLBI), other National Institute of Health (NIH) institutes and other foreign governmental and non-governmental organizations. The Discovery Phase analysis of sequence data is supported through UF1AG047133 (to Drs. Schellenberg, Farrer, Pericak-Vance, Mayeux, and Haines); U01AG049505 to Dr. Seshadri; U01AG049506 to Dr. Boerwinkle; U01AG049507 to Dr. Wijsman; and U01AG049508 to Dr. Goate and the Discovery Extension Phase analysis is supported through U01AG052411 to Dr. Goate, U01AG052410 to Dr. Pericak-Vance and U01 AG052409 to Drs. Seshadri and Fornage. Data generation and harmonization in the Follow-up Phases is supported by U54AG052427 (to Drs. Schellenberg and Wang).

The ADGC cohorts include: Adult Changes in Thought (ACT), the Alzheimer’s Disease Centers (ADC), the Chicago Health and Aging Project (CHAP), the Memory and Aging Project (MAP), Mayo Clinic (MAYO), Mayo Parkinson’s Disease controls, University of Miami, the Multi-Institutional Research in Alzheimer’s Genetic Epidemiology Study (MIRAGE), the National Cell Repository for Alzheimer’s Disease (NCRAD), the National Institute on Aging Late Onset Alzheimer’s Disease Family Study (NIA-LOAD), the Religious Orders Study (ROS), the Texas Alzheimer’s Research and Care Consortium (TARC), Vanderbilt University/Case Western Reserve University (VAN/CWRU), the Washington Heights-Inwood Columbia Aging Project (WHICAP) and the Washington University Sequencing Project (WUSP), the Columbia University Hispanic-Estudio Familiar de Influencia Genetica de Alzheimer (EFIGA), the University of Toronto (UT), and Genetic Differences (GD).

The CHARGE cohorts are supported in part by National Heart, Lung, and Blood Institute (NHLBI) infrastructure grant HL105756 (Psaty), RC2HL102419 (Boerwinkle) and the neurology working group is supported by the National Institute on Aging (NIA) R01 grant AG033193. The CHARGE cohorts participating in the ADSP include the following: Austrian Stroke Prevention Study (ASPS), ASPS-Family study, and the Prospective Dementia Registry-Austria (ASPS/PRODEM-Aus), the Atherosclerosis Risk in Communities (ARIC) Study, the Cardiovascular Health Study (CHS), the Erasmus Rucphen Family Study (ERF), the Framingham Heart Study (FHS), and the Rotterdam Study (RS). ASPS is funded by the Austrian Science Fond (FWF) grant number P20545-P05 and P13180 and the Medical University of Graz. The ASPS-Fam is funded by the Austrian Science Fund (FWF) project I904),the EU Joint Programme - Neurodegenerative Disease Research (JPND) in frame of the BRIDGET project (Austria, Ministry of Science) and the Medical University of Graz and the Steiermärkische Krankenanstalten Gesellschaft. PRODEM-Austria is supported by the Austrian Research Promotion agency (FFG) (Project No. 827462) and by the Austrian National Bank (Anniversary Fund, project 15435. ARIC research is carried out as a collaborative study supported by NHLBI contracts (HHSN268201100005C, HHSN268201100006C, HHSN268201100007C, HHSN268201100008C, HHSN268201100009C, HHSN268201100010C, HHSN268201100011C, and HHSN268201100012C). Neurocognitive data in ARIC is collected by U01 2U01HL096812, 2U01HL096814, 2U01HL096899, 2U01HL096902, 2U01HL096917 from the NIH (NHLBI, NINDS, NIA and NIDCD), and with previous brain MRI examinations funded by R01-HL70825 from the NHLBI. CHS research was supported by contracts HHSN268201200036C, HHSN268200800007C, N01HC55222, N01HC85079, N01HC85080, N01HC85081, N01HC85082, N01HC85083, N01HC85086, and grants U01HL080295 and U01HL130114 from the NHLBI with additional contribution from the National Institute of Neurological Disorders and Stroke (NINDS). Additional support was provided by R01AG023629, R01AG15928, and R01AG20098 from the NIA. FHS research is supported by NHLBI contracts N01-HC-25195 and HHSN268201500001I. This study was also supported by additional grants from the NIA (R01s AG054076, AG049607 and AG033040 and NINDS (R01 NS017950). The ERF study as a part of EUROSPAN (European Special Populations Research Network) was supported by European Commission FP6 STRP grant number 018947 (LSHG-CT-2006-01947) and also received funding from the European Community’s Seventh Framework Programme (FP7/2007-2013)/grant agreement HEALTH-F4-2007-201413 by the European Commission under the programme “Quality of Life and Management of the Living Resources” of 5th Framework Programme (no. QLG2-CT-2002-01254). High-throughput analysis of the ERF data was supported by a joint grant from the Netherlands Organization for Scientific Research and the Russian Foundation for Basic Research (NWO-RFBR 047.017.043). The Rotterdam Study is funded by Erasmus Medical Center and Erasmus University, Rotterdam, the Netherlands Organization for Health Research and Development (ZonMw), the Research Institute for Diseases in the Elderly (RIDE), the Ministry of Education, Culture and Science, the Ministry for Health, Welfare and Sports, the European Commission (DG XII), and the municipality of Rotterdam. Genetic data sets are also supported by the Netherlands Organization of Scientific Research NWO Investments (175.010.2005.011, 911-03-012), the Genetic Laboratory of the Department of Internal Medicine, Erasmus MC, the Research Institute for Diseases in the Elderly (014-93-015; RIDE2), and the Netherlands Genomics Initiative (NGI)/Netherlands Organization for Scientific Research (NWO) Netherlands Consortium for Healthy Aging (NCHA), project 050-060-810. All studies are grateful to their participants, faculty and staff. The content of these manuscripts is solely the responsibility of the authors and does not necessarily represent the official views of the National Institutes of Health or the U.S. Department of Health and Human Services.

The four LSACs are: the Human Genome Sequencing Center at the Baylor College of Medicine (U54 HG003273), the Broad Institute Genome Center (U54HG003067), The American Genome Center at the Uniformed Services University of the Health Sciences (U01AG057659), and the Washington University Genome Institute (U54HG003079).

Biological samples and associated phenotypic data used in primary data analyses were stored at Study Investigators institutions, and at the National Cell Repository for Alzheimer’s Disease (NCRAD, U24AG021886) at Indiana University funded by NIA. Associated Phenotypic Data used in primary and secondary data analyses were provided by Study Investigators, the NIA funded Alzheimer’s Disease Centers (ADCs), and the National Alzheimer’s Coordinating Center (NACC, U01AG016976) and the National Institute on Aging Genetics of Alzheimer’s Disease Data Storage Site (NIAGADS, U24AG041689) at the University of Pennsylvania, funded by NIA, and at the Database for Genotypes and Phenotypes (dbGaP) funded by NIH. This research was supported in part by the Intramural Research Program of the National Institutes of health, National Library of Medicine. Contributors to the Genetic Analysis Data included Study Investigators on projects that were individually funded by NIA, and other NIH institutes, and by private U.S. organizations, or foreign governmental or nongovernmental organizations.

The authors report no conflicts of interest.

We thank William J. Salerno for running Parliament.

