## Supplemental tables 1, 2, 4 and figure 1 for "A comparative study of structural variant calling strategies using the Alzheimer’s Disease Sequencing Project’s whole genome family data"

### Supplementary Tables

#### Supplementary Table 1

| Caller | File Size (GB) - Disc | File Size (GB) - Disc+ext | Avg. Run Time (hours) - Disc | Avg. Run Time (hours) - Disc+ext | Peak CPU % - Disc | Peak CPU % - Disc+ext | Peak Memory (GB) - Disc | Peak Memory (GB) - Disc+ext |
|---|---|---|---|---|---|---|---|---|
| Breakdancer | **209.05** [130.14-318.90] | **54.58** [75.32-37.29] | **4.94** [3.11-6.87] | **1.44** [0.77-2.14] | **22.70** [14.90-30.49] | **5.67** [3.73-7.62] | **23.26** [15.15-37.11] | **6.16** [5.23-9.00] |
| CNVnator | **209.05** [130.14-318.90] | **54.58** [75.32-37.29] | **1.79** [1.04-2.87] | **0.50** [0.40-0.69] | **4.43** [2.71-8.99] | **1.10** [0.78-1.60] | **2.23** [1.26-3.74] | **0.57** [0.49-0.82] |
| DELLY | **209.05** [130.14-318.90] | **54.58** [75.32-37.29] | **10.10** [6.60-16.64] | **2.78** [1.60-5.60] | **61.63** [37.58-80.95] | **16.61** [9.47-22.59] | **3.53** [2.05-5.71] | **0.96** [0.64 -1.65] |
| GATK | **209.05** [130.14-318.90] | **54.58** [75.32-37.29] | **5.86** [4.27-9.21] | **1.59** [1.10-2.48] | **76.73** [47.07-97.48] | **20.84** [11.40-29.88] | **1.35** [0.89-1.92] | **0.36** [0.23-0.49] |
| LUMPY | **209.05** [130.14-318.90] | **54.58** [75.32-37.29] | **13.33** [6.33-22.36] | **4.04** [2.87-5.79] | **13.34** [6.33-22.36] | **4.04** [2.89-5.79] | **5.92** [5.13-6.84] | **1.66** [0.80-2.61] |
| PINDEL | **209.05** [130.14-318.90] | **54.58** [75.32-37.29] | **18.13** [13.63-27.53] | **5.21** [3.41-7.46] | **20.59** [10.54-35.74] | **5.36** [3.98-6.93] | **9.39** [5.87-12.41] | **2.55** [1.45-3.24] |
| SWAN | **209.05** [130.14-318.90] | **54.58** [75.32-37.29] | **205.15** [155.39-262.87] | **57.10** [32.63-107.04] | **68.88** [47.14-77.27] | **19.00** [9.86-33.61] | **70.71** [32.14-108.91] | **18.41** [12.74-28.02 |

**Overview of computational performance for seven SV/indel callers**. The mean (bold), minimum, and maximum values are provided for each caller. Ten subjects were randomly selected from the discovery (Disc) and discovery-extension (Disc+ext) phases. The second and third columns provide the file sizes in gigabytes. The fourth and fifth columns provide the average runtime in hours. The sixth and seventh columns provide the peak CPU usage. Columns eight and nine provide the peak memory in gigabytes.

| Supplementary Table 2. Candidate AD Genes |
| --- |
| *ABCA7* |
| *ABCG1* |
| *ABI3* |
| *ACE* |
| *ADAM10* |
| *ADAMTS1* |
| *AKAP9* |
| *APOE* |
| *APP* |
| *BIN1* |
| *BZRAP1* |
| *CASP7* |
| *CASS4* |
| *CD2AP* |
| *CD33* |
| *CELF1* |
| *CLU* |
| *COBL* |
| *CR1* |
| *CTDP1* |
| *ECHDC3* |
| *EPHA1* |
| *FBXL7* |
| *FERMT2* |
| *FRMD4A* |
| *GALNT7* |
| *GAS2L2* |
| *GCH1* |
| *GLIS1* |
| *GLIS3* |
| *GPAA* |
| *HBEGF* |
| *HDAC9* |
| *IGHV1-67* |
| *INPP5D* |
| *IQCK* |
| *KANSL1* |
| *KCNJ15* |
| *KCNMB2* |

| |
|---|
| *MAPT* |
| *MEF2C* |
| *MS4A4A* |
| *MS4A6A* |
| *NCR2* |
| *NME8* |
| *NOTCH3* |
| *OPRL1* |
| *OSBPL6* |
| *OSTN* |
| *PCDH8* |
| *PDCL3* |
| *PICALM* |
| *PILRA* |
| *PLCG2* |
| *PLD3* |
| *PLXNA4* |
| *PSEN1* |
| *PSEN2* |
| *PTK2B* |
| *PTPRG* |
| *SERPINB1* |
| *SHARPIN* |
| *SLC10A2* |
| *SLC24A4* |
| *SORL1* |
| *TM2D3* |
| *TP53INP1* |
| *TPBG* |
| *TREM2* |
| *TRIP4* |
| *TTC3* |
| *UNC5C* |
| *ZCWPW1* |
| *ZNF655* |

**Supplementary Table 4. Deletion Size Frequency Distribution**

| Bin Size | Scalpel | | Parliament | | Comments |
|---|---|---|---|---|---|
| 20-99 | 82,180 | 88.69% | 2,253 | 10.13% | |
| 100-199 | 3,984 | 4.30% | 6,601 | 29.67% | |
| 200-299 | 1,502 | 1.62% | 2,455 | 11.04% | |
| 300-399 | 2,745 | 2.96% | 3,195 | 14.36% | ALU Peak |
| 400-499 | 662 | 0.71% | 982 | 4.41% | |
| 500-599 | 518 | 0.56% | 730 | 3.28% | |
| 600-699 | 429 | 0.46% | 529 | 2.38% | |
| 700-799 | 376 | 0.41% | 359 | 1.61% | |
| 800-899 | 242 | 0.26% | 257 | 1.16% | |
| 900-999 | 21 | 0.02% | 221 | 0.99% | |
| 1,000-1,999 | NA | | 1,321 | 5.94% | SVA Peak |
| 2,000-2,999 | NA | | 855 | 3.84% | |
| 3,000-3,999 | NA | | 572 | 2.57% | |
| 4,000-4,999 | NA | | 403 | 1.81% | |
| 5,000-9,999 | NA | | 983 | 4.42% | L1 Peak |
| 10,000-99,999 | NA | | 530 | 2.38% | |
| Total* | 92,659 | 100% | 22,246 | 100% | |

* Singletons excluded

### Supplementary Tables – Large

Supplementary Table 3.  All deletions - characteristics

Supplementary Table 5.  Deletions near AD genes.

Supplementary Table 6.  Co-segregation of confirmed deletions near AD genes in multiplex families.

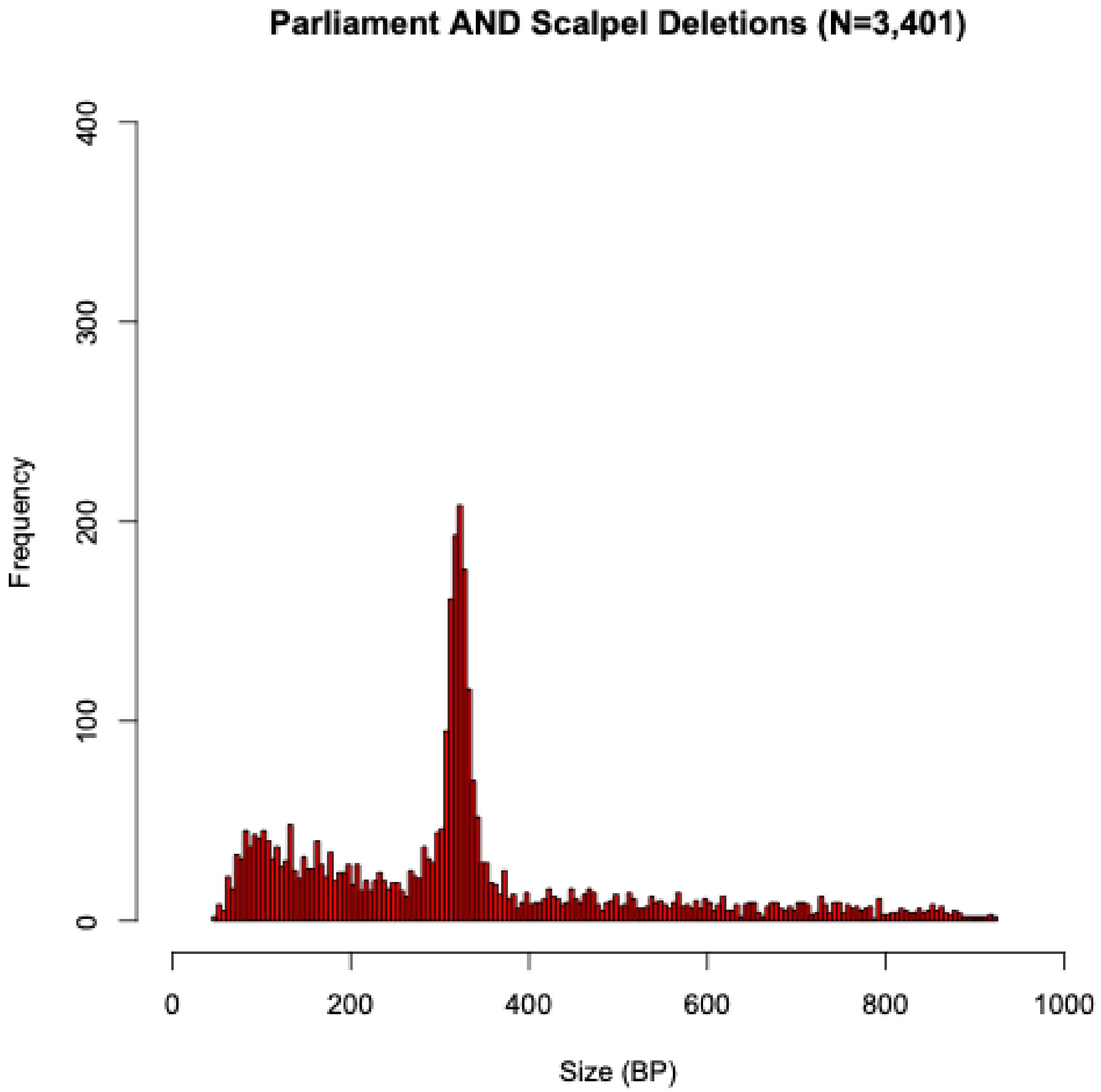


**Supplementary Figure 1.**
